# OxyR senses reactive sulfane sulfur and activates genes for its removal in *Escherichia coli*

**DOI:** 10.1101/561019

**Authors:** Ningke Hou, Zhenzhen Yan, Kaili Fan, Huanjie Li, Rui Zhao, Yongzhen Xia, Huaiwei Liu, Luying Xun

## Abstract

Reactive sulfane sulfur species such as hydrogen polysulfide and organic persulfide are newly recognized as normal cellular components, involved in signaling and protecting cells from oxidative stress. Their production is extensively studied, but their removal is less characterized. Herein, we showed that reactive sulfane sulfur is toxic at high levels, and it is mainly removed via reduction by thioredoxin and glutaredoxin with the release of H_2_S in *Escherichia coli*. OxyR is best known to respond to H_2_O_2_, and it also played an important role in responding to reactive sulfane sulfur under both aerobic and anaerobic conditions. It was modified by hydrogen polysulfide to OxyR C199-SSH, which activated the expression of thioredoxin 2 and glutaredoxin 1. This is a new type of OxyR modification. Bioinformatics analysis showed that OxyRs are widely present in bacteria, including strict anaerobic bacteria. Thus, the OxyR sensing of reactive sulfane sulfur may represent a conserved mechanism for bacteria to deal with sulfane sulfur stress.

## Introduction

H_2_S has been proposed as a gasotransmitter because it is involved in many physiological and pathological processes in animals and plants, such as ageing *(Hine et al, 2015)*, neuromodulation *(Abe & Kimura, 1996)*, cancer cell proliferation *(Cai et al, 2010)*, metabolic reprogramming *(Gao et al, 2015)*, and stomatal closure in plant *(Lisjak et al, 2010)*. The mechanism of H_2_S signaling is often via protein persulfidation or S-sulfhydration. Since H_2_S cannot direct react with protein thiols, its oxidation product reactive sulfane sulfur, which can readily react with thiols to generate persulfides, has been identified *(Mishanina et al, 2015; Toohey, 2011)*. Reactive sulfane sulfur species include hydrogen polysulfides (H_2_S_n_, n≥2), organic polysulfides (RSS_n_H, RSS_n_R, n≥2), and organic persulfides (RSSH), which can also be produced directly from cysteine or cystine, and they are now considered as normal components in both prokaryotic and eukaryotic cells *(Sawa et al, 2018; Yadav et al, 2016)*. They possess both nucleophilic and electrophilic characteristics, while thiols (cysteine, GSH, etc.) are generally nucleophilic *(Ono et al, 2014; Park et al, 2015)*. As nucleophiles, they are better reductants than thiols *(Ida et al, 2014)*; as electrophiles, the electrophilic sulfane sulfur (S^0^) can be transferred to protein thiols to generate protein-SSH, which modifies enzyme activities and protects proteins from irreversible oxidation *(Mustafa et al, 2009; Paul & Snyder, 2015)*. Owing to these unique dual-reactivities, reactive sulfane sulfur is involved in many cellular processes, such as redox homeostasis maintenance, virulence regulation in pathogenic bacteria, and biogenesis of mitochondria *(Fujii et al, 2018; Peng et al, 2017)*. Albeit the good roles, sulfane sulfur may be toxic at high concentrations. Indeed, elemental sulfur has been used as an antimicrobial agent for ages, and its efficiency is likely impaired by its low solubility *(Williams & Cooper, 2010)*. Due to its low solubility, elemental sulfur is not considered as a reactive sulfane sulfur species. Advances in the synthesis of sulfur nanoparticles have significantly increased the antimicrobial efficiency of sulfur *(Rai et al, 2016)*. Sulfur is often used as a fungicide. Although its toxicity mechanism is unclear, a recent study suggested that sulfur is transported into the cell in the form of hydrogen polysulfide *(Sato et al, 2011)*, inducing protein persulfidation as a possible toxic mechanism *(Islamov et al, 2018)*. Fungi may use glutathione to reduce polysulfides to H_2_S as a detoxification mechanism *(Samrat et al, 2013; Sato et al, 2011)*. Organosulfur compounds can be used to treat antibiotic-resistant bacteria, and they are converted to hydrogen polysulfide inside the cells for the toxicity *(Xu et al, 2018)*. Both bacteria and fungi show reduced viability being exposed to sulfane sulfur stress *(Sato et al, 2011; Xu et al, 2018)*. Therefore, intracellular sulfane sulfur is likely maintained within a range for microorganisms under normal conditions.

Multiple pathways for sulfane sulfur generation have been discovered. Cystathionine β-synthase and cystathionine γ-lyase produce sulfane sulfur from cystine *(Ida et al, 2014)*; 3-mercaptopyruvate sulfurtransferase (3-MST) and cysteinyl-tRNA synthetase (CARS) produce sulfane sulfur from cysteine *(Akaike et al, 2017; Nagahara et al, 2018)*; Sulfide:quinone oxidoreductase and superoxide dismutase (SOD) produce sulfane sulfur from H_2_S *(Olson et al, 2018; Xin et al, 2016)*. Catalase can oxidize H_2_S and polysulfides to sulfur oxides *(Olson et al, 2017)*. Most microorganisms possess several of these pathways. On the flipside, elimination pathways are less investigated. Aerobic microorganisms may apply persulfide dioxygenase to remove excessive sulfane sulfur *(Xia et al, 2017)*, and the persulfide dioxygenase expression can be induced by sulfane sulfur via sulfane sulfur-sensing transcription factors *(H et al, 2017; Lira et al, 2018; Luebke et al, 2014)*.

For anaerobic microorganisms that dominate in the intestinal tract, their sulfane sulfur elimination pathways are ambiguous. A reasonable hypothesis is that they use glutathione (GSH) or NADPH to reduce sulfane sulfur to H_2_S and then release it out of cells *(Carbonero et al, 2012)*, as observed in anaerobically cultured fungi *(Abe et al, 2007; Sato et al, 2011)*. A recent report that two thioredoxin-like proteins catalyze the reduction of protein persulfidation in *Staphylococcus aureus* also support the hypothesis *(Peng et al, 2017)*. However, it is unclear whether sulfane sulfur induces the expression of these enzymes.

*Escherichia coli*, a common intestinal bacterium, contains three thioredoxins and four glutaredoxins. The expression of TrxA, GrxB, and GrxD is regulated by guanosine 3’,5’-tetraphosphate, and the expression of GrxC is regulated by cAMP, both of which are nutrient related messengers *(Lim et al, 2000; Srivatsan & Wang, 2008)*. These four enzymes are highly abundant in *E. coli*; together they account for more than 1% of total protein *(Fernandes et al, 2005; Fernandes & Holmgren, 2004; Ritz et al, 2000)*. The expression of GrxA, TrxC, and KatG (catalase) is regulated by OxyR upon exposure to H_2_O_2_. Without oxidative stress, these proteins are much lower than other thioredoxins and glutaredoxins *(Fernandes & Holmgren, 2004; Gutierrez-Ríos et al, 2007; Ritz et al, 2000)*.

OxyR was initially identified as a regulator responding to reactive oxygen species (ROS) *(Christman et al, 1985; Storz et al, 1990)*. ROS triggers the formation of intra disulfide bond between Cys^199^ and Cys^208^ or oxidizes Cys^199^ to C199-SOH. The exact mechanism is still in debate *(Choi et al, 2001; Kim et al, 2002; Zheng et al, 1998)*. Three additional modifications on Cys199 (C199-SNO, C199-SSG and avicinylation) are also known, which result in different OxyR configurations, DNA binding affinities, and promoter activities *(Haridas et al, 2005; Kim et al, 2002; Seth & Stamler, 2012)*. Thus, OxyR leads to multi-levels of transcriptional responses when responding to different stress signals.

Herein, we systematically investigated the sulfane sulfur reduction activity and expression pattern of thioredoxins, glutaredoxins, and KatG in *E. coli*. We found that these enzymes are responsible for maintaining the homeostasis of intracellular sulfane sulfur. Further investigation unveiled a new and H_2_S_n_-specific modification of OxyR.

## Results

### The accumulation and reduction of endogenous sulfane sulfur in E. coli

*E. coli* has endogenous sulfane sulfur producing enzymes, including CARS, 3-MST, SodA, and SodC *(Akaike et al, 2017; Nagahara et al, 2018; Olson et al, 2018)*. We cultured early-log phased *E. coli* cells in LB medium and used the sulfane sulfur sensitive probe SSP4 (Fig. 1A) to detect intracellular sulfane sulfur. The intracellular sulfane sulfur started to accumulate at middle-log phase and reached the maximum at early stationary phase. On the other hand, when the stationary-phased cells were transferred into fresh medium (OD_600_=1), their intracellular sulfane sulfur decreased quickly with concomitant release of H_2_S (Fig. 1B). This phenomenon suggests that sulfane sulfur may be reduced to H_2_S by enzymes, such as thioredoxin and glutaredoxin *(Dóka et al, 2016; Wedmann et al, 2016)*.

**Fig. 1.**
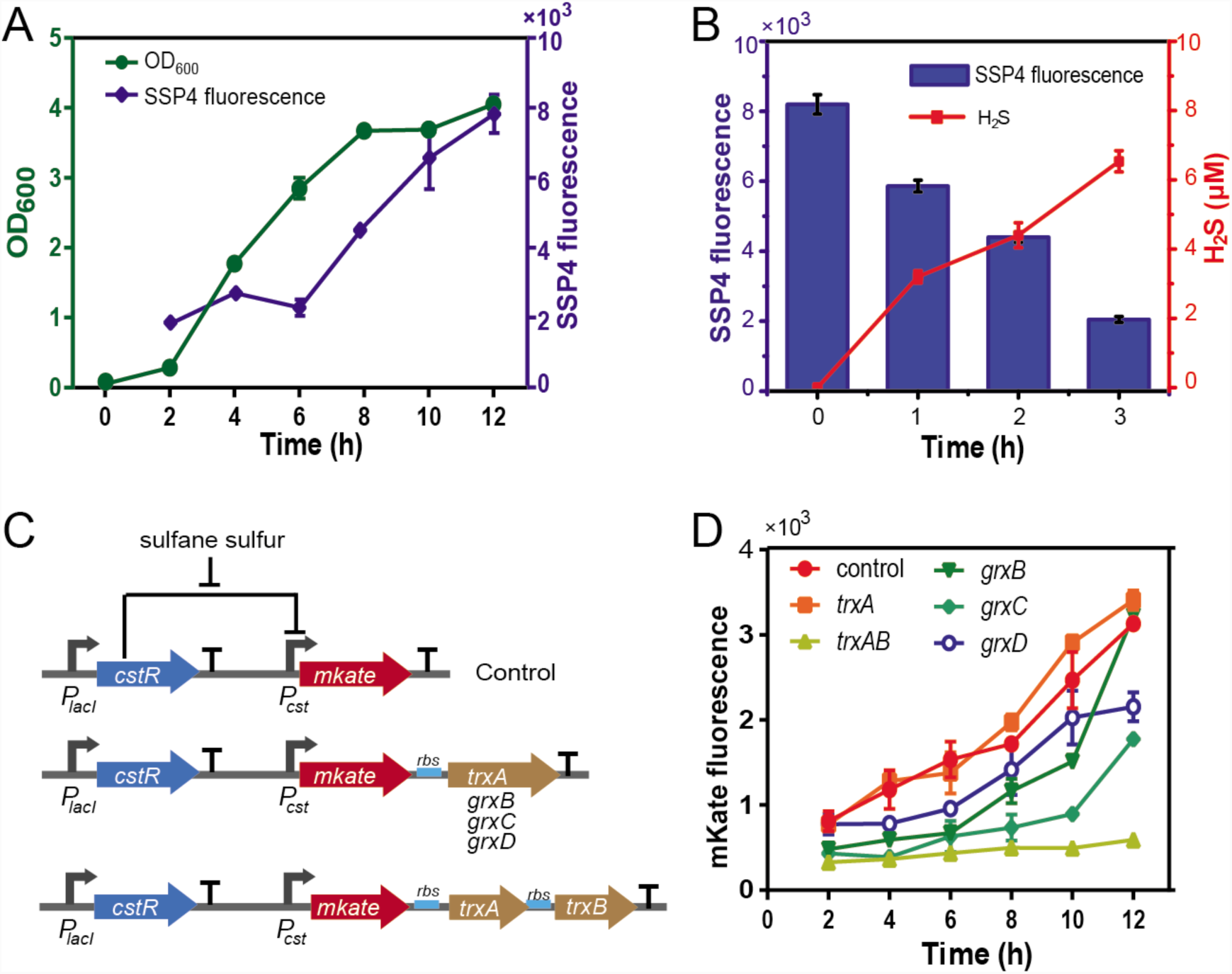
Endogenous sulfane sulfur production and reduction in *E. coli*. A) *E. coli* accumulated sulfane sulfur and released H_2_S during late-log and stationary phases. B) Stationary-phased *E. coli* reduced intracellular sulfane sulfur to H_2_S after being transferred to fresh LB. C) CstR-based reporters for real-timely monitoring intracellular sulfane sulfur. D) Overexpression of thioredoxins and glutaredoxins decreases intracellular sulfane sulfur, as indicated by mKate fluorescence.

### Thioredoxin and glutaredoxin participate in the reduction of intracellular sulfane sulfur

To confirm the change of intracellular sulfane sulfur *in vivo*, we constructed a transcription factor (TF)-based reporting plasmid, which contained a sulfane sulfur-sensing TF (CstR) *(Luebke et al, 2014)*, its cognate promoter (*Pcst*), and a red fluorescent protein (mKate, with a C-terminus degradation tag ssrA) (Fig. 1C). Using the reporting plasmid, the increase of intracellular sulfane sulfur in live cells (Fig. 1A) was reported as the mKate fluorescence (Fig. 1D). When GrxB, GrxC, or GrxD was co-transcribed with mKate under the control of CstR, their expression could partially decrease the sulfane sulfur accumulation as reflected with the mKate fluorescence intensity (Fig. 1D). When TrxA was co-transcribed, it did not affect sulfane sulfur accumulation (Fig. 1D). However, when thioredoxin reductase (TrxB) was co-expressed with TrxA, sulfane sulfur were not increased during the log phase of growth (Fig. 1D). These results indicate that thioredoxin and glutaredoxin reduce sulfane sulfur inside *E. coli*.

The artificial operons contained a negative feedback loop when coupled with an enzyme that reduces sulfane sulfur (Fig. 1C&D). The loop effectively maintained intracellular sulfane sulfur levels within a narrow range, defined by the leaky strength of *Pcst* and the sensitivity of CstR as well as the controlled enzyme activity. Since OxyR is known to regulate similar enzymes, such as TrxC, GrxA, and KatG, we speculated that OxyR may play a role similar as CstR in the artificial operons (Fig. 1C).

### OxyR alleviates sulfane sulfur stress by regulating the expression of TrxC, GrxA, and KatG under both aerobic and anaerobic conditions

We deleted *oxyR* gene in *E. coli* and observed that the mutant became more sensitive to exogenously added H_2_S_n_ under both aerobic or anaerobic conditions (Fig. 2A). After complementing *oxyR* into *ΔoxyR*, the strain reassumed the tolerance to H_2_S_n_ to the same level of the wild type (*wt*) (Fig. 2A). The results indicated that OxyR play an important role in dealing with the exogenously sulfane sulfur stress both under aerobic or anaerobic conditions. Compared with *wt, ΔoxyR* had higher intracellular sulfane sulfur at log-phase (Fig. 2B). In addition, when the *ΔoxyR* cells at the stationary phase were transferred into fresh LB medium at OD_600_ of 1, the decrease of the intracellular sulfane sulfur and release of H_2_S were slower than that of the *wt* cells (Fig. 2C and Fig. 1B). The results suggested that OxyR responds to sulfane sulfur and activates the expression of sulfane sulfur reduction enzymes.

**Fig. 2.**
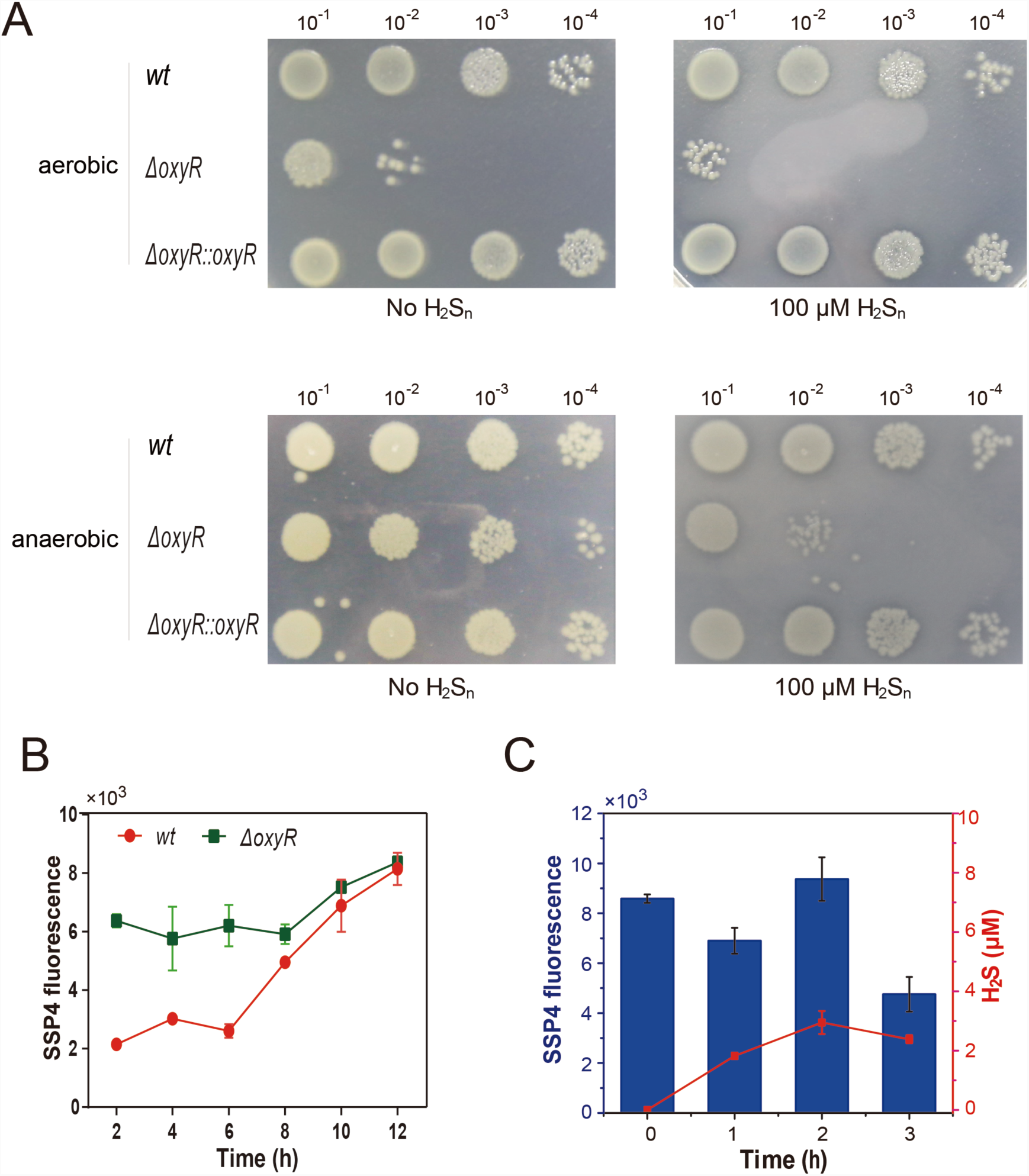
OxyR affects RSS reduction in *E. coli*. A) *E. coli ΔoxyR* is more sensitive to exogenous H_2_S_n_ stress. B) *E. coli ΔoxyR* accumulates more endogenous sulfane sulfur than *E. coli wt* during growth in LB. C) Stationary-phased *E. coli ΔoxyR* reduces endogenous sulfane sulfur to H_2_S more slowly than *E. coli wt* (Fig. 1B) after being transferred to fresh LB.

To confirm OxyR responds to H_2_S_n_ and activates the expression of *trxC, grxA* and *katG*, we constructed three reporting plasmids with an *mKate* gene under the control of a *trxC, grxA*, or *katG* promoter. These plasmids were transformed into *E. coli wt* and *ΔoxyR*. The recombinant cells were subjected to H_2_S_n_ stress under aerobic conditions. In *wt*, all three promoters led to a low *mKate* expression in the absence of H_2_S_n_ but resulted in obviously higher expression when H_2_S_n_ was added (Fig. 3A). Whereas in the *ΔoxyR* strain, the three promoters led to constantly low expression of *mKate* with or without H_2_S_n_ stress (Fig. 3B). After introducing *oxyR* back to *ΔoxyR*, the promoters performed the same as that in *wt* (Fig. 3C). Further, overexpression of *trxC, grxA*, and *katG* in *E. coli ΔoxyR* decreases intracellular sulfane sulfur (Fig. S1). The induction by H_2_S_n_ was further conformed by *in vitro* transcription-translation experiments. DTT or H_2_S_n_ treated-OxyR and a DNA fragment containing the *trxC* promoter and *mKate* (*P_trxC_-mKate*) were added into the cell-free transcription-translation system. When DTT-treated OxyR (the reduced form) was used, *mKate* expression was low. Whereas, when H_2_S_n_-treated OxyR was used, *mKate* expression was significantly increased (Fig. 4). These results indicated that H_2_S_n_ induces the *trxC* promoter via directly modifying OxyR.

**Fig. 3.**
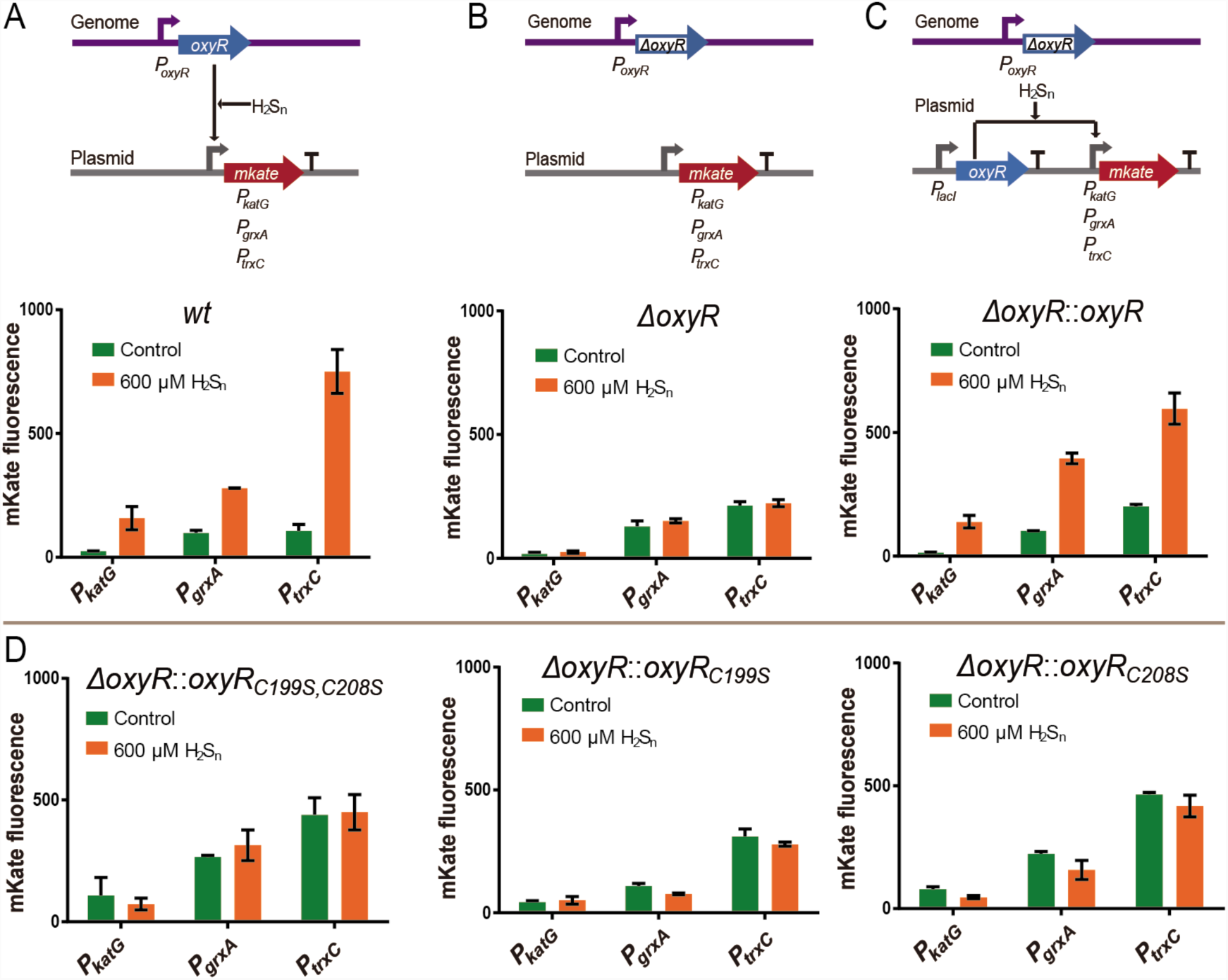
H_2_S_n_ upregulates expression of *katG, grxA*, and *trxC* via OxyR under aerobic conditions. A) H_2_S_n_ induces expression of *katG, grxA*, and *trxC* in *E. coli wt*. B) The induction effect is lost in *E. coli ΔoxyR.* C) OxyR complementation recovers the induction effect. D) Cys199 and Cys208 single or double mutants lost the induction effect.

**Fig. 4.**
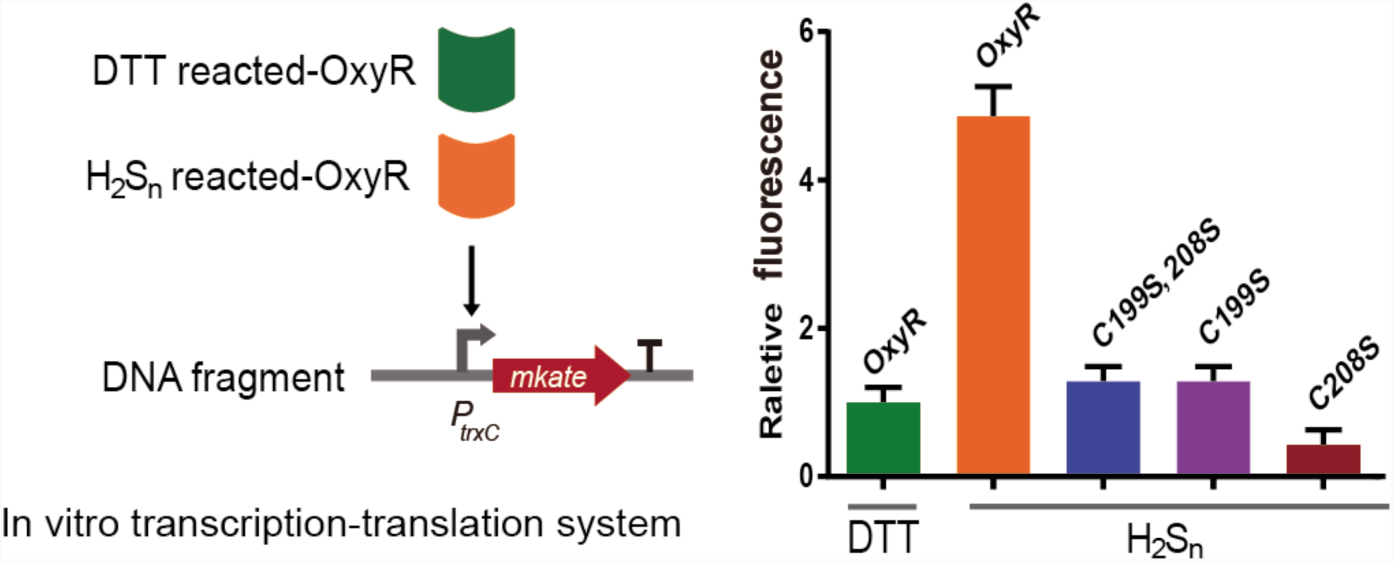
*In vitro* transcription-translation analysis of H_2_S_n_ activation of OxyR and its mutants.

We also test the induction by H_2_S_n_ under anoxic conditions using qPCR as mKate does not mature under anaerobic conditions. Similarly, *katG, grxA*, and *trxC* had higher expression in *wt* when 200 μM H_2_S_n_ were added (Fig. 5A), but not in *ΔoxyR* (Fig. 5B). After complementing *oxyR* to *ΔoxyR*, the induction was resumed (Fig. 5C).

**Fig. 5.**
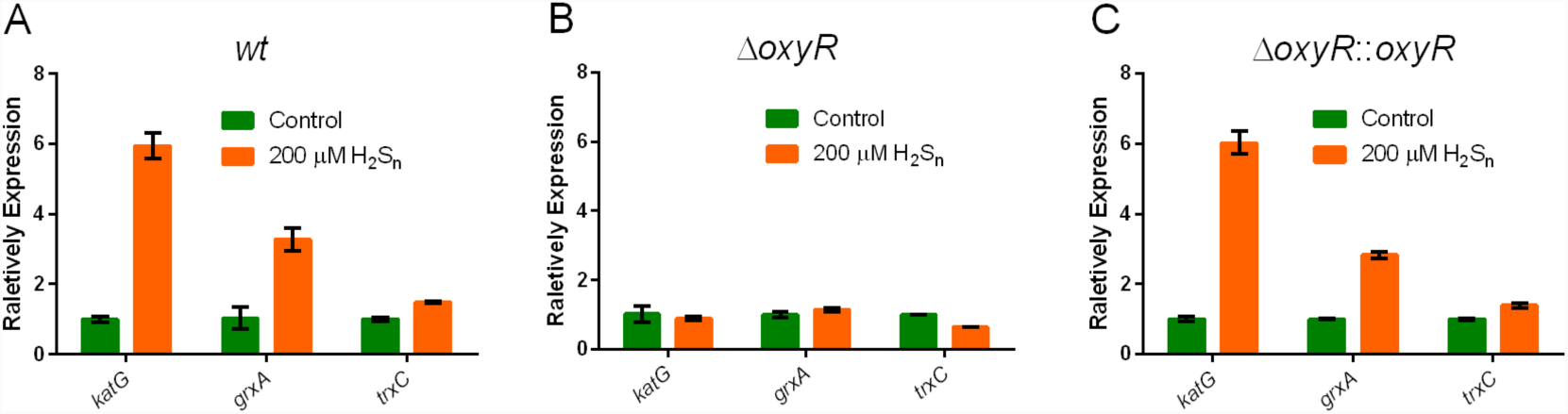
H_2_S_n_ upregulates expression of *katG, grxA*, and *trxC* via OxyR under anoxic conditions. A) H_2_S_n_ induces expression of *katG, grxA*, and *trxC* in *E. coli wt*. B) The induction effect is lost in *E. coli ΔoxyR.* C) OxyR complementation recovers the induction effect.

Since the H_2_S_n_ solution contained some sulfide, we tested if sulfide could induce the gene expression. Sulfide did not induce the expression of related genes in *wt* (Fig. 6B), excluding the signal function of sulfide. When we used *E. coli* cells harboring a sulfide:quinone oxidoreductase (SQR) of *C. pinatubonensis* JMP134, the added sulfide was oxidized to H_2_S_n_ *(Xin et al, 2016)*, which induced the expression of *trxC, grxA* and *katG* (Fig. 6A).

**Fig. 6.**
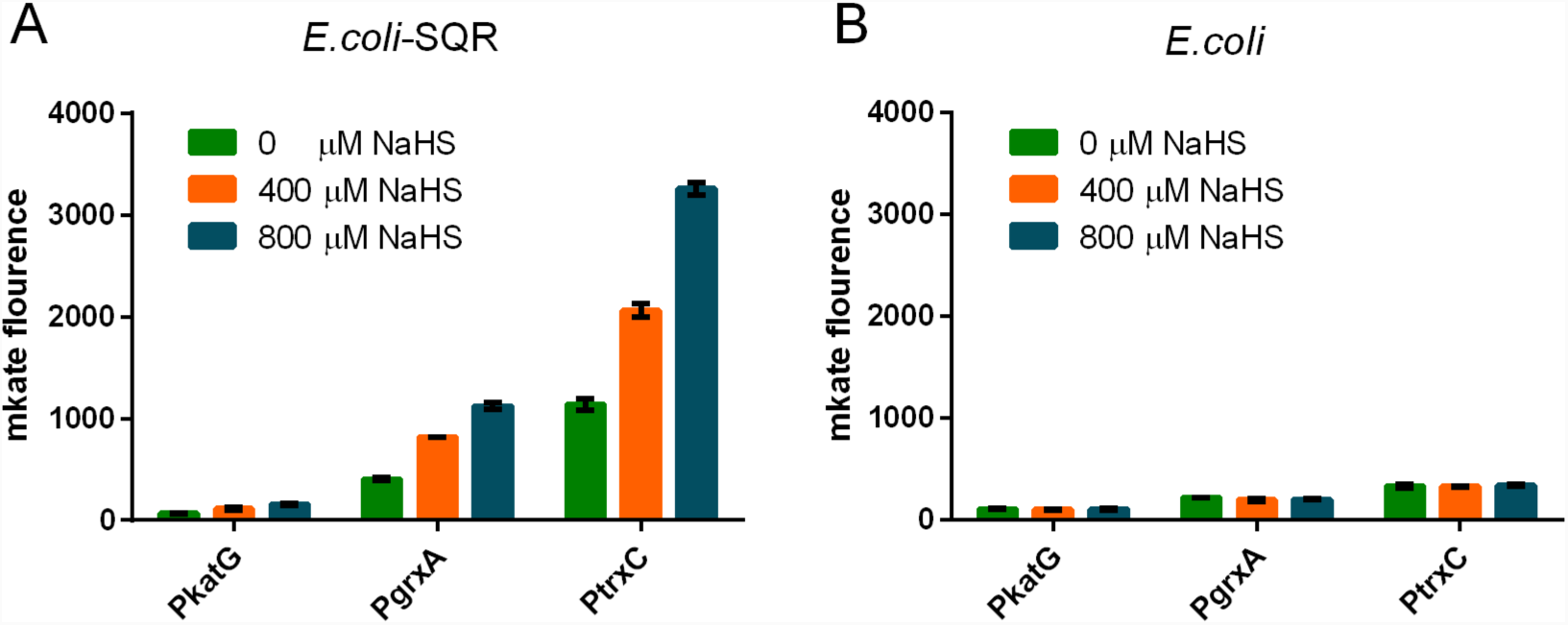
The induction of *trxC, grxA and katG* by NaHS with (A) or without (B) SQR. The *sqr* of *C. pinatubonensis* JMP134 gene was expressed under the *P_lacI_* promoter in the pBBrMCS2 plasmid.

### The molecular mechanism of OxyR sensing H_2_S_n_

OxyR contains six cysteine residues. Previous studies indicated two of them (Cys^199^ and Cys^208^) are involved in ROS sensing *(Zheng et al, 1998)*. We constructed an OxyR_4C→A_ mutant (except for Cys^199^ and Cys^208^, the other four cysteines were mutated to alanines) and expressed it in *ΔoxyR*. The mutant regulated *trxC, grxA*, or *katG* promoters essentially the same as the wild-type OxyR in the presence of H_2_S_n_. Whereas, OxyR_C199S_, OxyR_C208S_, and OxyR_C199S; C208S_ all lost the regulation function (Fig. 3D). Together, these results indicated that the same as in ROS sensing, only Cys^199^ and Cys^208^ are involved in H_2_S_n_ sensing.

To find out the molecular mechanism on how OxyR senses H_2_S_n_, mass spectrometry analysis was performed to analyze the H_2_S_n_-treated OxyR. A short peptide (MW: 1356.67) containing Cys^199^ but not Cys^208^ was identified (peptide 1, Fig. 7 and Fig. S2) and about 20% of it contained a persulfidation on Cys^199^ (MW: 1388.64) (peptide 2, Fig. 7 and Fig. S3), according to the peak area in MS^1^ spectrogram. A peptide containing Cys^208^ was also found, but the Cys^208^ was unmodified by iodoacetamide (IAM) (MW: 2144.87) (peptide 3, Fig. 7 and Fig. S4). Cys^208^ was not modified by IAM indicating that it is not accessible to IAM, consistent with a previous report that Cys^208^ is buried in the protein *(Kim et al, 2002)*. No peptide containing both Cys^199^ and Cys^208^ was detected. These data collectively indicated that OxyR senses H_2_S_n_ via persulfidation on Cys^199^, other than forming disulfide or –S_n_– (n≥3) bond between Cys^199^ and Cys^208^.

**Fig. 7.**
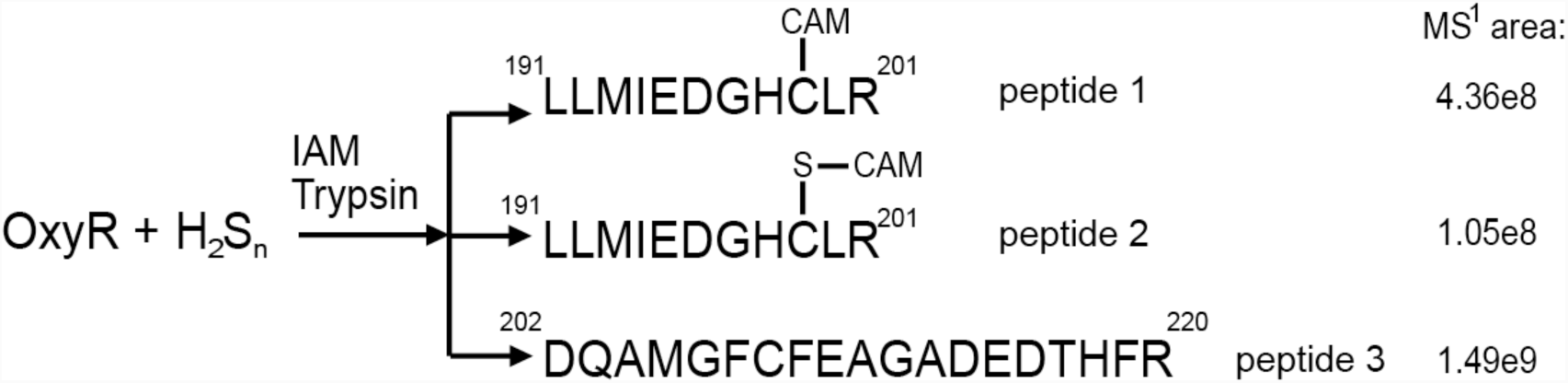
LTQ-Orbitrap tandem mass analysis of H_2_S_n_-reacted OxyR. MS data of the peptides are provided in Fig. S2∼4.

### Global transcriptome analysis of H_2_S_n_-stressed and H_2_O_2_-stressed E. coli

To systematic understand the effect of H_2_S_n_ on *E. coli* and any similarities with the H_2_O_2_ stress, we analyzed transcriptome response of *E. coli* with or without H_2_S_n_ stress. In the presence of 400 μM H_2_S_n_, *E coli* significantly downregulated its energy metabolism-related genes, such as maltoporin (71.3 fold-change) and glycerol-3-phosphate transporter (33.7 fold-change). The most up regulated genes included phage holin (425.9 fold-change), phage recombination protein Bet (96 fold-change), and a putative single-stranded DNA binding protein, suggesting that H_2_S_n_ is toxic to *E. coli (Xu et al, 2018)*. When we checked sulfane sulfur-removing enzymes, GrxA had the highest (2.1 fold) and TrxC and KatG had mild (1.6 fold and 1.5 fold) transcriptional increases in H_2_S_n_-stressed cells (Fig. 8A), consistent with the results of fluorescence reporting systems (Fig. 3); whereas, the other sulfane sulfur-reducing enzymes TrxA, TrxB, and GrxD had no obvious transcriptional change, and GrxB and GrxC had a slight decrease (Fig. 8A and 8B). These proteins are regulated by nutrient related regulators and are highly abundant in *E. coli (Lim et al, 2000; Srivatsan & Wang, 2008) (Fernandes et al, 2005; Fernandes & Holmgren, 2004; Ritz et al, 2000)*, suggesting that they may play the “house-keeping” role and the OxyR related GrxA, TrxC and KatG are in stress responses..

**Fig. 8.**
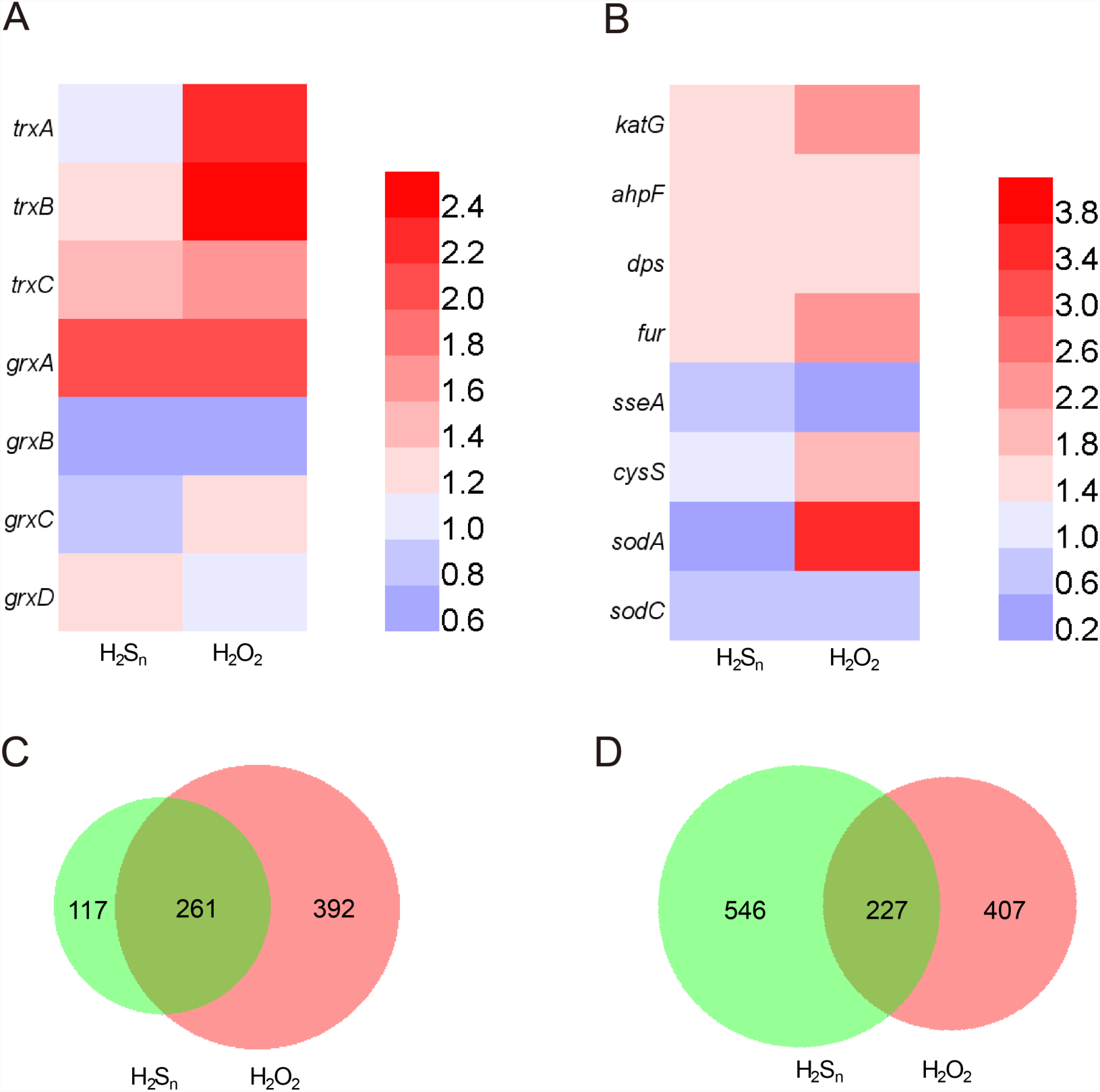
Transcriptome analysis of H_2_S_n_- or H_2_O_2_-stressed *E. coli*. A, B) Transcriptional change of genes involved in sulfane sulfur production and reduction. C) Numbers of transcriptionally upregulated genes by H_2_S_n_ and H_2_O_2_ stresses. D) Numbers of transcriptionally downregulated genes by H_2_S_n_ and H_2_O_2_ stresses.

**Fig. 9.**
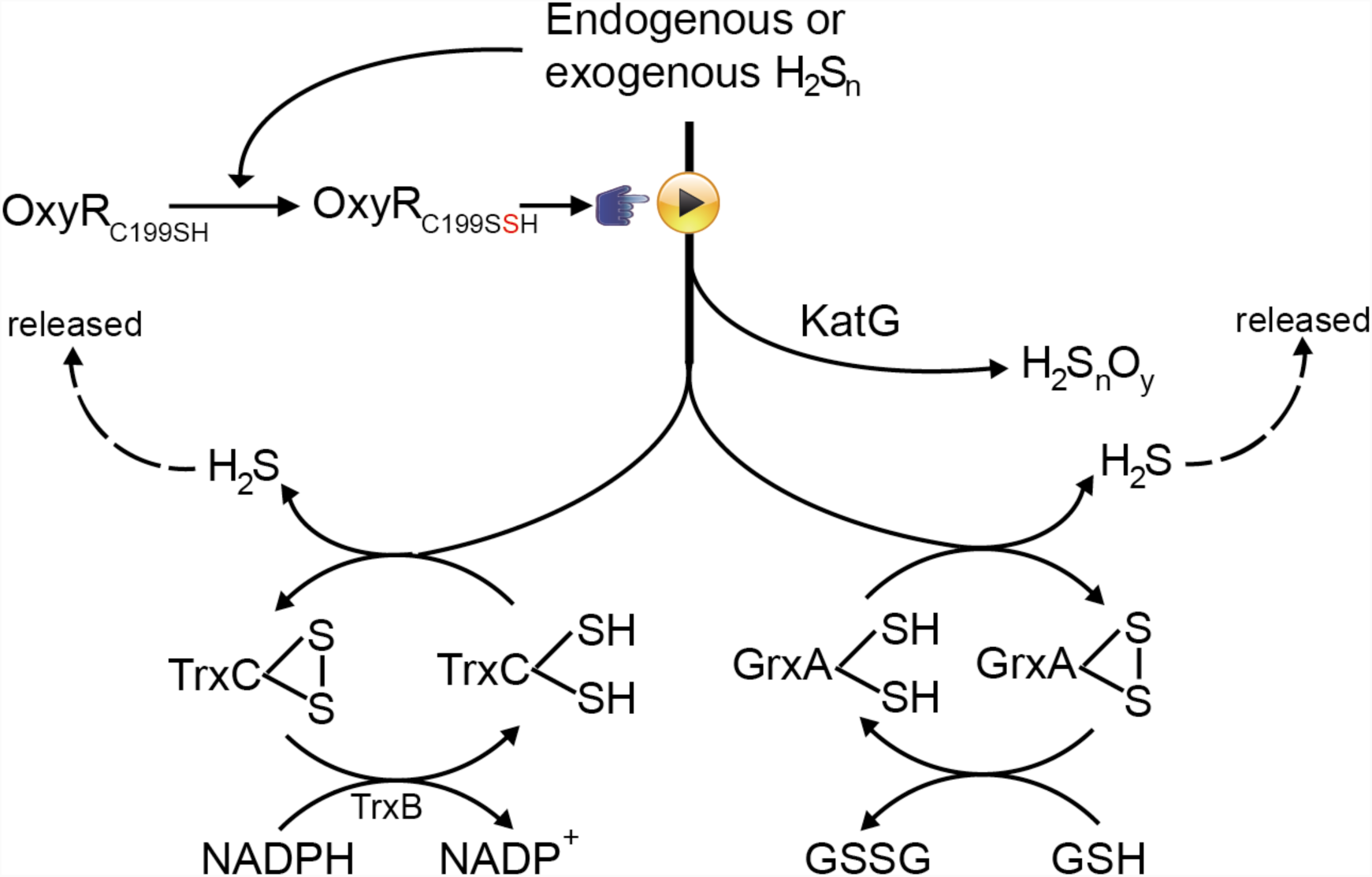
Schematic representation of the OxyR-regulated RSS reduction pathways in *E. coli*.

At a global level, there were also similarities and differences. Among the 1286 genes affected by H_2_O_2_, upregulated and downregulated ones were 652 and 634, respectively; whereas, among the 1150 genes affected by H_2_S_n_ stress, only about 1/3 were upregulated (Fig. 8C and D). Gene ontology (GO) analysis indicated the cellular processes affected by them were different. For instance, H_2_S_n_ stress upregulated more genes pertaining to cellular component, e.g., cell part (GO:0044464) and macromolecular complex (GO:0032991), and downregulated more genes pertaining to molecular function, such as molecular transducer activity (GO:0060089) and signal transducer activity (GO:0004871); whereas H_2_O_2_ stress upregulated more genes pertaining to molecular function, e.g., ribonucleotide binding (GO:0032553) and carbohydrate derivative binding (GO:0097367), and downregulated no gene pertaining to cellular component (Fig. S5 and S6). The TCA cycle is upregulated by H_2_S_n_ stress but downregulated by H_2_O_2_ stress; biosynthesis of secondary metabolites (i.e. serine hydroxymethyltransferase, beta-gulcosidase, 3-deoxy-7-phosphoheptulonate synthase, etc.) is downregulated by H_2_S_n_ stress but not affected by H_2_O_2_ stress (Fig. S7 and S8).

H_2_S_n_ stress downregulated the expression of 3-MST (encoded by sseA), SodA, and SodC, but did not affect the expression of CARS (encoded by cysS) (Fig. 8B). These four enzymes are all involved in sulfane sulfur generation. Whereas, H_2_O_2_ stress significantly upregulated the expression of CARS and SodA, consistent with the report that proteins involved in sulfane sulfur biosynthesis are induced under oxidative stress because sulfane sulfur functions as antioxidants *(Fukuto et al, 2018)*. GrxA, TrxC, and KatG were similarly upregulated by H_2_S_n_ and H_2_O_2_ stress; Trx A and TrxB were not obviously affected by H_2_S_n_ stress, but H_2_O_2_ stress upregulated them (Fig. 8A and B). Overall, the transcriptomic data indicated that H_2_S_n_ and H_2_O_2_ stresses lead to largely different responses in *E. coli*. However, some proteins like TrxC, GrxA, and KatG, are likely involved in alleviating both stresses.

### The distribution of OxyR in sequenced bacterial genomes

Several sulfane sulfur-sensing TFs have been discovered, and all are involved in regulating the genes involved in sulfur oxidation *(H et al, 2017; Lira et al, 2018; Luebke et al, 2014)*. OxyR is the first global gene regulator that also senses sulfane sulfur. Thus, we invested the distribution of OxyR among 8286 microbial genomic sequences (NCBI updated until November 11, 2017) by using BLAST search, and then confirmed with the conserved domain and phylogenetic tree analysis. 4772 identified OxyR distributed in 4494 bacterial genomes, including 2432 Gammaproteobacteria, 887 Bataproteobacteria, 478 Alphaproteobacteria, 287 Corynebacteriales, 130 Flavobacteiriia, 67 Streptomycetales, and 63 Bacterioidia; the other 24 classes had few genomes containing OxyR (Table S1). Thus, OxyR is widely distributed in bacteria. It is worth noting that OxyR is also present in many obligate anaerobic bacteria, such as *Bacteroides* spp., *Prevotella* spp., and *Porphromonas* spp. Some of them are common residents in the human gut. For anaerobic bacteria, OxyR is more likely to deal with H_2_S_n_ stress than H_2_O_2_ stress, as the latter is not an issue under anaerobic conditions. It is noteworthy that intestinal tract is an anoxic and sulfur rich environment *(Daeffler et al, 2017; Espey, 2013)*. Hence for intestinal bacteria such as *E. coli* and *Salmonella enterica*, OxyR may play important roles for their survival in the gut.

## Discussion

In this study, we reported the fifth type modification of OxyR, C199-SSH, which is modified by H_2_S_n_. H_2_S_n_-modified OxyR upregulates the expression of TrxC, GrxA, and KatG; these enzymes can convert sulfane sulfur to releasable H_2_S or H_2_S_n_O_y_ (Fig. 9). Other OxyR-regulated proteins like hydroperoxide reductase AhpF, DNA protection protein Dps, and transcriptional regulator Fur are also upregulated under H_2_S_n_ stress (Fig. 8A). Therefore, as other modifications *(Haridas et al, 2005; Kim et al, 2002; Seth & Stamler, 2012)*, C199-SSH should also lead to multi-level transcriptional responses. Although Cys^208^ is not modified by H_2_S_n_, it plays a critical role during H_2_S_n_ sensing, probably stabilizing the active C199-SSH.

Reactive sulfane sulfur species are essential intracellular contents. They are beneficial at low levels *(Ida et al, 2014; Mustafa et al, 2009; Paul & Snyder, 2015)*; however, they are toxic at high levels. Our systematic study of H_2_S_n_ stress unveiled that TrxA, TrxB, GrxB, GrxC and GrxD may function as a house-keeping machinery to prevent the buildup of intracellular RRS to toxic levels; OxyR-regulated GrxA, TrxC and KatG may function as emergency backups to deal environmentally confronted or abnormally over-accumulated sulfane sulfur. Glutathione redoxins and thioredoxins reduce sulfane sulfur to H_2_S that is released out of the cell for microorganisms growing under anaerobic conditions *(Abe et al, 2007; Sato et al, 2011; Xia et al, 2017)*. For bacteria or animal host with SQR, the released H_2_S is captured and oxidized back to sulfane sulfur under aerobic conditions *(Lagoutte et al, 2010)*. For *E. coli* and bacteria without SQR, H_2_S will be released even under aerobic conditions *(Li et al, 2019; Xia et al, 2017)*.

S and O are both chalcogens. Reactive sulfane sulfur species are similar chemicals to ROS (e.g., HSSH *vs* H_2_O_2_) *(Deleon et al, 2016)*; their modifications to proteins are also analogous, i.e., protein-SSH vs protein-SOH *(Mishanina et al, 2015)*. From an evolutionary perspective, the former’s history can be traced back before the Great Oxidation Event (GOE), when O_2_ had not been generated by cyanobacteria. As an abundant element on ancient earth, S should play important roles in ancient microorganisms. Therefore, sulfur metabolism related enzymes had emerged before the oxygen’s era. It is reasonable to speculate that the anti-ROS proteins are derived from anti-sulfane sulfur ones *(Olson et al, 2017)*. Possibly that is the reason why the anti-ROS network largely overlaps with that of anti-sulfane sulfur. On the other hand, we also observed that *E. coli* has obviously responding-discrepancies when confronting H_2_O_2_ or HSSH (Fig. 7). These discrepancies are in agreement with the multi-level transcriptional responses of OxyR when activated by different reagents *(Haridas et al, 2005; Kim et al, 2002; Seth & Stamler, 2012)*.

In conclusion, we discovered that *E. coli* uses thioredoxins, glutaredoxins, and catalase to control homeostasis of intracellular sulfane sulfur. OxyR functions as a reactive sulfane sulfur sensor via persulfidation of its Cys^199^ both under aerobic or anoxic conditions. This is the fifth type modification observed for OxyR activation. Since OxyR is widely distributed in both aerobic and anaerobic bacteria, the OxyR-regulated network may represent a conserved mechanism that bacteria can resort to when confronting endogenous and/or exogenous sulfane sulfur stress.

## Materials and Methods

### Strains, plasmids, and chemicals

All strains and plasmids used in this study are listed in Table S2. Deletion of *oxyR* was performed following a reported method *(Datsenko & Wanner, 2000)*. *E. coli* strains were grown in Lysogeny broth (LB) medium. Antibiotics (50 μg/ml) were added when required. SSP4 (3’,6’-Di(*O*-thiosalicyl)fluorecein) was purchased from DOJINDO MOLECULAR TECHNOLOGIES *(Bibli et al, 2018)*. H_2_S_n_ was prepared by following Kamyshny & Alexey’s method *(Kamyshny et al, 2009)*. Briefly, 13 mg of sulfur powder and 70 mg of sodium sulfide were added to 5 ml of anoxic distilled water under argon gas. The pH was adjusted to 9.3 with 6 M HCl. The obtained product contained a mixture of H_2_S_n_, where n varies from 2 to 8 *(Olson et al, 2017)*, but at low concentration and neutral pH, H_2_S_2_ is dominant *(Bogdándi et al, 2018; Xin et al, 2016)*.

### Endogenous sulfane sulfur analysis

SSP4 probe was used for batch analysis. Cells were washed with and resuspended in HEPES buffer (50 mM, pH 7.4); then 10 μM SSP4 and 0.5 mM CTAB were added. After an incubation at 37°C for 15 min in the dark with gently shaking (125 rpm), reagents were washed off with HEPES buffer (50 mM, pH 7.4). Reacted-cells were subjected to flow cytometry (FACS) analysis by using BD Accuri™ C5. For each sample, >10,000 cells were analyzed in FL1-A channel. The average fluorescent intensity was used.

The CstR-based reporting system was used for real-time analysis. *cstR* gene was chemically synthesized by Genewiz (Shanghai) company and expressed with *P_lacI_* promoter in pTrcHis2A plasmid, where the *trc* promoter was replaced by the CstR cognate promoter, and a *mkate* gene (with a C-terminus degradation tag *ssrA*) was put after it (Table S2, entry 22). For *trxA, trxB, grxB, grxC*, or *grxD* overexpression experiment, the gene was put after *mkate*, separated by an *rbs* sequence (Table S2, entries 23∼27). *E. coli* strains containing reporting plasmids were culture in LB medium at 37°C with shaking (220 rpm). Fluorescence was analyzed by FACS (FL3-A channel, >10,000 cells).

### Hydrogen sulfide production analysis

Production of hydrogen sulfide was determined using a previously reported method *(Kimura et al, 2015)*. Briefly, hydrogen sulfide was derivatized with mBBr then analyzed by HPLC (LC-20A, Shimadzu) equipped with a fluorescence detector (RF-10AXL, Shimadzu). A C18 reverse phase HPLC column (VP-ODS, 150×4 mm, Shimadzu) was pre-equilibrated with 80% Solvent A (10% methanol and 0.25% acetic acid) and 20% Solvent B (90% methanol and 0.25% acetic acid). The column was eluted with the following gradients of Solvent B: 20% from 0 to 10 min; 20%–40% from 10 to 25 min; 40%–90% from 25 to 30 min; 90%–100% from 30 to 32 min; 100% from 32 to 35 min; 100 to 20% from 35 to 37 min; and 20% from 37 to 40 min. The flow rate was 0.75 ml/min. For detection, the excitation wavelength was set to 340 nm and emission wavelength was set to 450 nm.

### H_2_S_n_ inhibition and induction tests

For growth inhibition test, middle-log phased *E. coli* cells (OD600=0.8) were diluted and dripped in freshly prepared LB agar medium containing 0 or 100 μM H_2_S_n_ and incubated in 37°C under aerobic conditions. For anaerobic conditions, the anaerobic LB agar plates were prepared in an anaerobic glove box and the dilution and drip of *E. coli* cells also performed in an anaerobic glove box, then incubated in an anaerobic incubator at 37°C for 24 hours. For promoter induction test, a *mkate* gene was put after *trxC, grxA*, or *katG* native promoter in pTrchis2A plasmid (Table S2, entries 17∼19). The *oxyR* or its mutant gene was expressed under the *P_lacI_* promoter in the same plasmid (Table S2, entries 5∼16) for complementary experiments. The obtained plasmids were transformed into *wt* and *ΔoxyR* strains. Early log-phased *E. coli* cells (OD_600_= 0.5, in liquid LB) were incubated with 600 μM H_2_S_n_ for 2 hours. Cells were harvested and washed with HEPES buffer (50 mM, pH 7.4), then subjected to FACS analysis (FL3-A channel, >10,000 cells).

### Real-time quantitative reverse transcription PCR (RT-qPCR)

RNA sample was prepared by using the TRIzol™ RNA Purification Kit (12183555, Invitrogen). Total cDNA was synthesized using the All-In-One RT Master Mix (ABM). For RT-qPCR, strains were grown in anaerobic LB medium until OD_600_ reached 0.4, and then 200 μM H_2_S_n_ were added into anaerobic bottle. After 60 min, cells were collected by centrifugation and RNA was extracted. RT-qPCR was performed by using the Bestar SybrGreen qPCR Mastermix (DBI) and LightCycler 480II (Roche). For calculation the relative expression levels of tested genes, GAPDH gene expression was used as the internal standard.

### Protein purification and reaction with DTT or H_2_S_n_

The *oxyR* gene with a C-terminal His tag was ligated into pET30. Mutants of *oxyR* were constructed from this plasmid via site-directed mutagenesis *(Xia et al, 2015)*. The obtained plasmids were transformed into *E. coli* BL21 (DE3). For protein expression, *E. coli* cells were cultured in LB medium at 25°C with shaking (150 rpm) until OD_600_ reacted 0.6–0.8, 0.4 mM isopropyl-β-D-thiogalactopyranoside (IPTG) was added, and cells were cultured for additional 16 hours at 16°C. Cells were then harvested and disrupted through a high pressure cracker SOCH-18 (STANSTED); protein was purified via the Ni-NTA resin (Invitrogen). Buffer exchange of the purified protein was performed by using PD-10 desalting column (GE Healthcare).

Reactions were performed in an anaerobic glove box. 0.6 mg/ml protein was mixed with 200 mM DTT in a pH 8.0 buffer (50 mM NaH_2_PO_4_, 300 mM NaCl). After 1-hour incubation at RT, the protein was dialyzed against 0.5 M KCl until the dialysis buffer was free of DTNB-titratable SH group. For H_2_S_n_ reaction, the reduced OxyR was mixed with about 10-fold concentration of H_2_S_n_ and incubated for 30 min at RT. Unreacted H_2_S_n_ was removed via dialysis. The reacted-proteins were sealed and taken out from the glove box to be used in further experiments.

### LC-MS/MS analysis of OxyR

The H_2_S_n_-reacted OxyR (0.5 mg/ml) was mixed with iodoacetamide (IAM), and then digested with trypsin by following a previously reported protocol *(H et al, 2017)*. The Prominence nano-LC system (Shimadzu) equipped with a custom-made silica column (75 μm × 15 cm) packed with 3-μm Reprosil-Pur 120 C18-AQ was used for the analysis. For the elution process, a 100 min gradient from 0% to 100% of solvent B (0.1 % formic acid in 98% acetonitrile) at 300 nl/min was used; solvent A was 0.1 % formic acid in 2% acetonitrile. The eluent was ionized and electrosprayed via LTQ-Orbitrap Velos Pro CID mass spectrometer (Thermo Scientific), which run in data-dependent acquisition mode with Xcalibur 2.2.0 software (Thermo Scientific). Full-scan MS spectra (from 400 to 1800 m/z) were detected in the Orbitrap with a resolution of 60,000 at 400 m/z.

### In vitro transcription-translation analysis

*In vitro* translation-transcription reactions were performed using the Purexpress *In Vitro* Protein Synthesis system (NEB #E6800). The reaction solution was prepared in the following order: 10 μL solution A (NEB #E6800), 7.5 μL solution B (NEB #E6800), 2 μL *E. coli* RNA polymerase (NEB #M0551), 1 μL RNase inhibitor, 500 ng DTT- or H_2_S_n_- reacted protein, 200 ng DNA fragment containing P*_trxC_*-mKate, and RNase free water. The total volume was 25 μL. The solution was incubated at 37°C for 3 hours. After reaction, the translated mKate was diluted four times with distilled water, and assayed by using the Synergy H1 microplate reader. The excitation wavelength was set to 588 nm, and the emission wavelength was set to 633 nm. The fluorescence intensity from reduced OxyR was used as standard; fluorescence intensities from other groups were divided by the standard to calculate the relative expression levels.

### Transcriptomic analysis

*E. coli wt* strain was cultured in LB medium until OD_600_ reached 0.5, and 500 μM H_2_S_n_ or 500 μM H_2_O_2_ were added. After 20 min of treatment, cells were harvested and total RNA was extracted by using the TRIzol™ RNA Purification Kit (12183555,Invitrogen). RNA quality was assessed with the RNA Nano 6000 Assay Kit of the Agilent Bioanalyzer 2100 system (Agilent Technologies). rRNA was removed with the Ribo-Zero rRNA Removal Kit (MRZMB 126, Epicentre Biotechnologies). For cDNA library construction, first-strand cDNA was synthesized by using random hexamer primers from fragmentation of mRNA and second-strand cDNA was synthesized by using a dNTP mixture containing dUTP with DNA polymerase I and RNase H. After adenylation of the ends of blunt-ended DNA fragments, NEBNext index adaptor oligonucleotides were ligated to the cDNA fragments. The second-strand cDNA containing dUTP was digested with the USER enzyme. The first-strand DNA fragments with ligated adaptors on both ends were selectively enriched in a 10-cycle PCR reaction, purified (AMPure XP), and the library was quantified using the Agilent High Sensitivity DNA assay on the Agilent Bioanalyzer 2100 system. The library was sequencing on Illumina Hiseq 2500 platform. Sequencing was performed at Beijing Novogene Bioinformatics Technology Co., Ltd. The clean data were obtained from raw data by removing reads containing adapter, poly-N and low quality reads. The clean reads were aligned with the genome of *E. coli* BL21 by using Bowtie2-2.2.3. Gene expression was quantified as reads per kilobase of coding sequence per million reads (RPKM) algorithm. Genes with a p-value<0.05 found by DESeq and change fold>1.5 were considered as significantly differentially expressed. Gene Ontology (GO) and KEGG analyses were performed at NovoMagic platform provided by Beijing Novogene Bioinformatics Technology Co., Ltd.

### Analysis of OxyR distribution in sequenced bacterial genomes

A microbial genomic protein sequence set from NCBI updated until November 11, 2017 was downloaded for OxyR search. The query sequences of OxyR were reported OxyR proteins *(Choi et al, 2001; Inseong et al, 2015; Kaewkanya et al, 2003)* and were used to search the database by using Srandalone BLASTP algorithm with conventional criteria (e-value ≤ 1e^-5^, coverage ≥ 45%, identity ≥ 30%) to obtain OxyR candidates from 8286 bacterial genomes. A conserved domain PBP2_OxyR and PRK11151 were used as standard features for further filtration of OxyR candidates. The candidates containing PBP2_OxyR or PRK11151 were identified as putative OxyR.

## Acknowledgments

The work was financially supported by grants from the National Natural Science Foundation of China (91751207, 31770093), the National Key Research and Development Program of China (2016YFA0601103), and the Natural Science Foundation of Shandong Province (ZR2016CM03, ZR2017ZB0210).

