## Supplementary material for "OxyR senses reactive sulfane sulfur and activates genes for its removal in *Escherichia coli*": Supplmental Figures 1-8 and Tables 1-2

**Figure S1-8, Table S1-2**

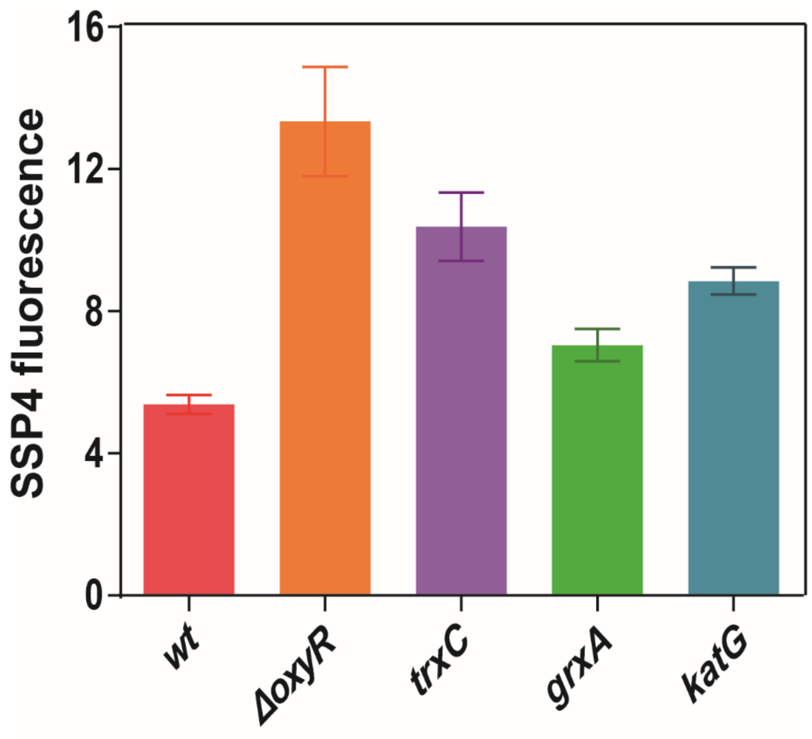


**Figure S1. Overexpression of *trxC*, *grxA*, and *katG* in *E. coli ΔoxyR* decreases intracellular reactive sulfane sulfur.** *TrxC*, *grxA*, and *katG* genes were expressed with *PlacI* promoter in pTrcHis2A plasmids. Cells were cultured in LB medium until $OD_{600}$ reached 2 and then measured the endogenous reactive sulfane sulfur using SSP4.

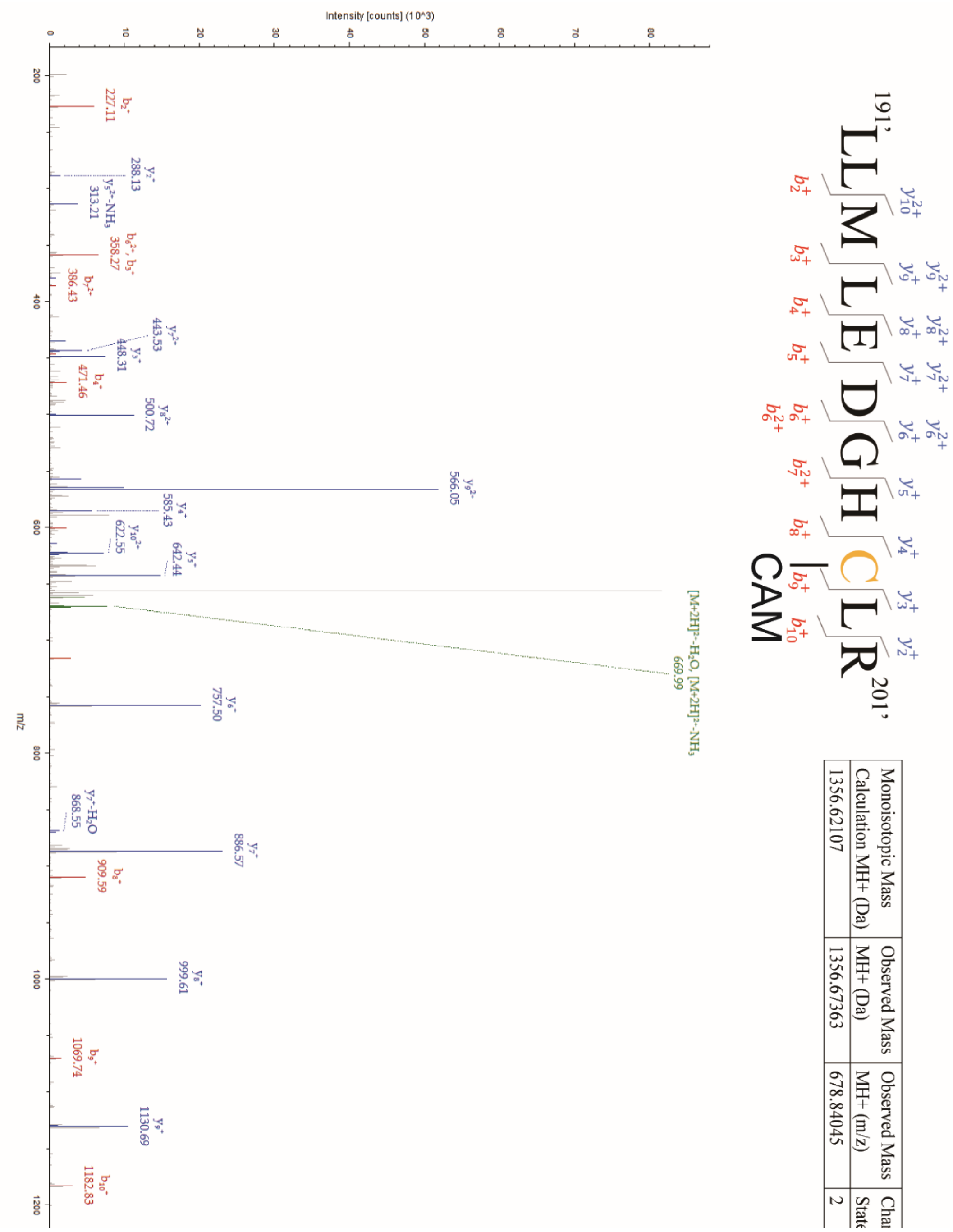


**Figure S2. MS$^2$ data of peptide 1 of HSSH-reacted OxyR.** The sample was digested by trypsin and analyzed by LTQ-Orbitrap Tandem MS. The –SH group was blocked by IAM in the peptide fragment containing $Cys_{199}$.

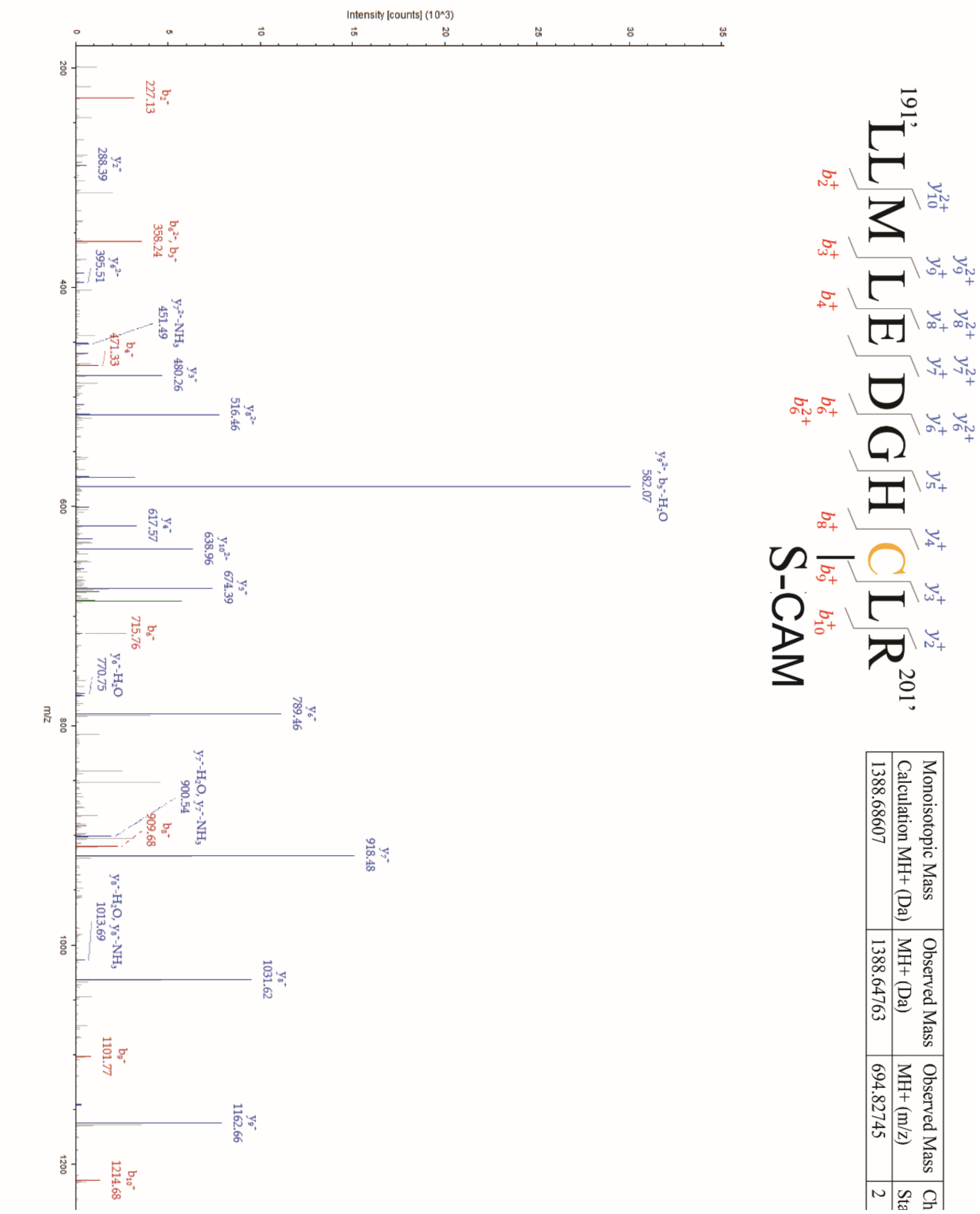


| Monoisotopic Mass Calculation MH+ (Da) | Observed Mass MH+ (Da) | Observed Mass MH+ (m/z) | Ch Sta |
|---|---|---|---|
| 1388.68607 | 1388.64763 | 694.82745 | 2 |

**Figure S3. MS$^2$ data of peptide 2 of HSSH-reacted OxyR.** The sample was digested by trypsin and analyzed by LTQ-Orbitrap Tandem MS. The –SSH group was blocked by IAM in the peptide fragment containing $Cys_{199}$.

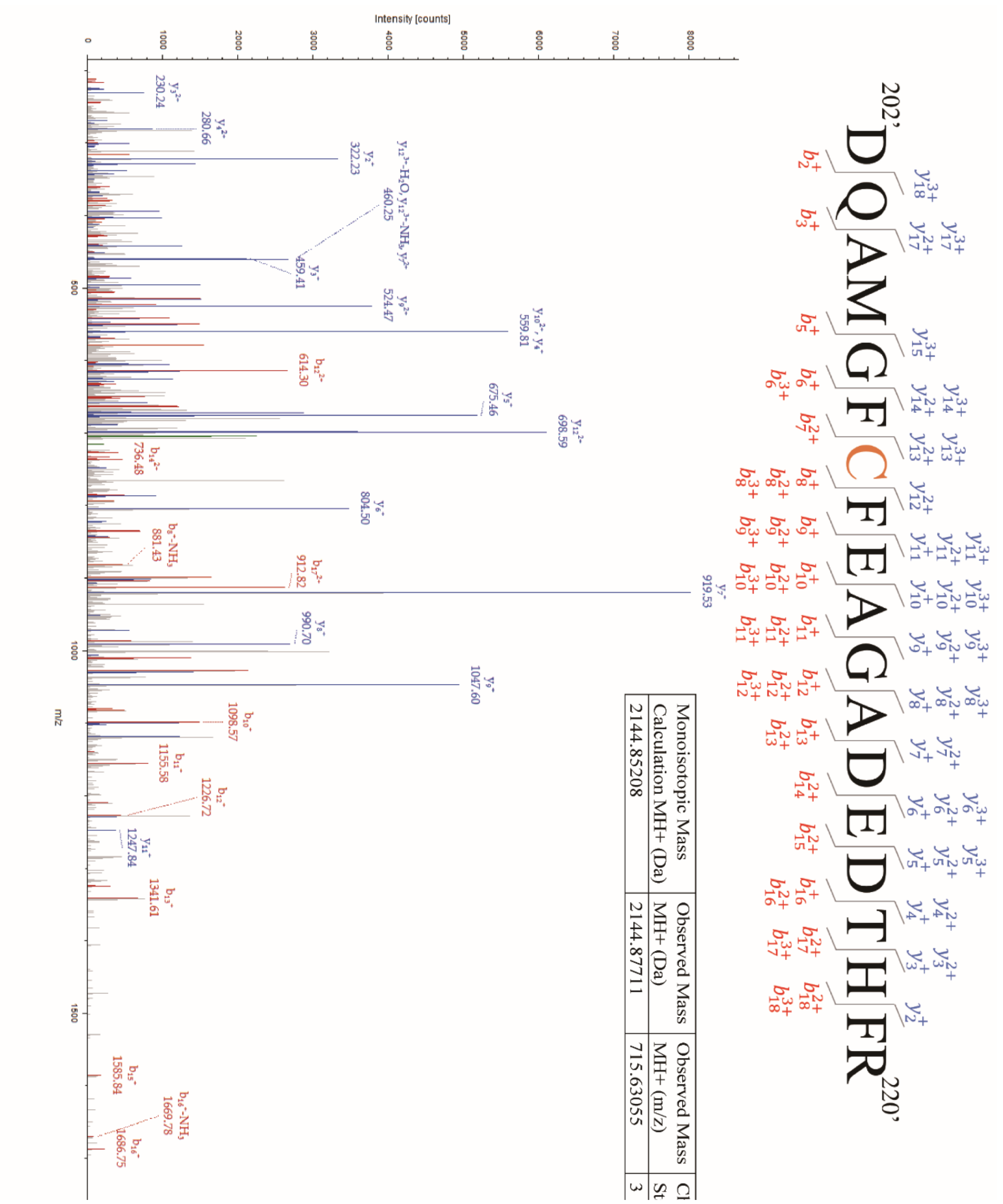


**Figure S4. MS$^2$ data of peptide 3 of HSSH-reacted OxyR.** The sample was digested by trypsin and analyzed by LTQ-Orbitrap Tandem MS. $Cys_{208}$ contained –SH group in the peptide fragment.

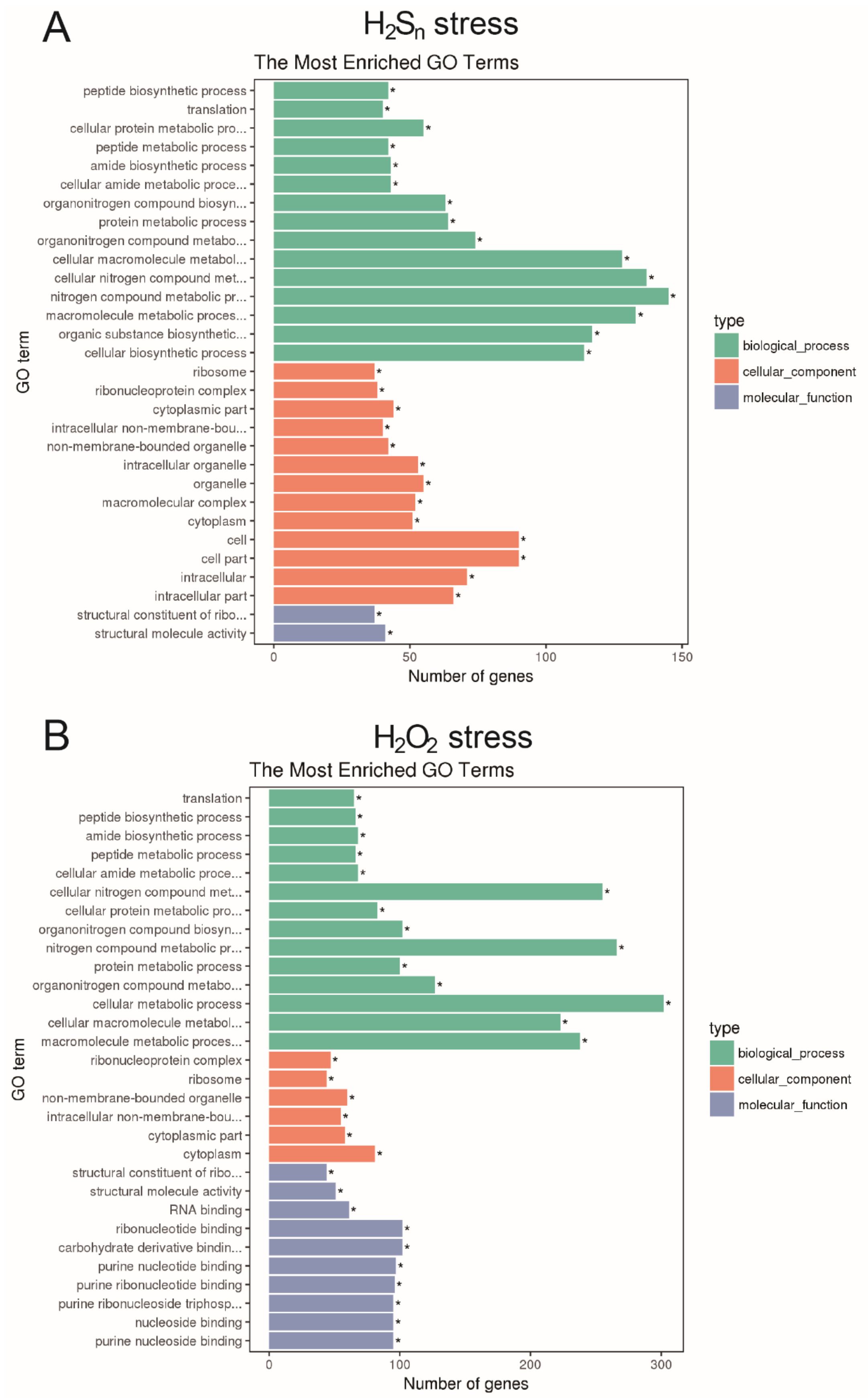


**Figure S5. Gene ontology classification of upregulated genes in $H_2S_n$ and $H_2O_2$ stressed *E. coli*.**

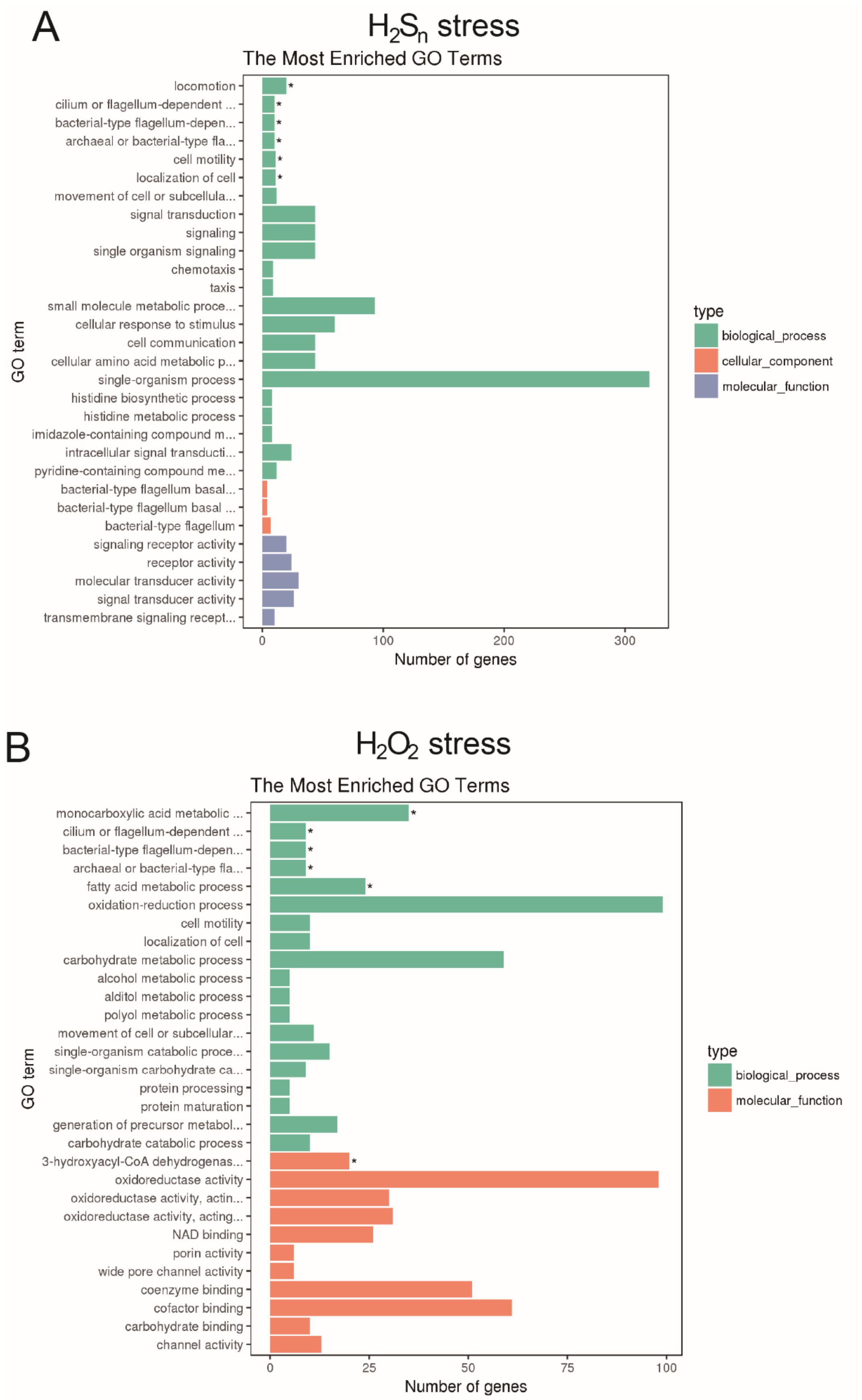


**Figure S6. Gene ontology classification of downregulated genes in $H_2S_n$ and $H_2O_2$ stressed *E. coli*.**

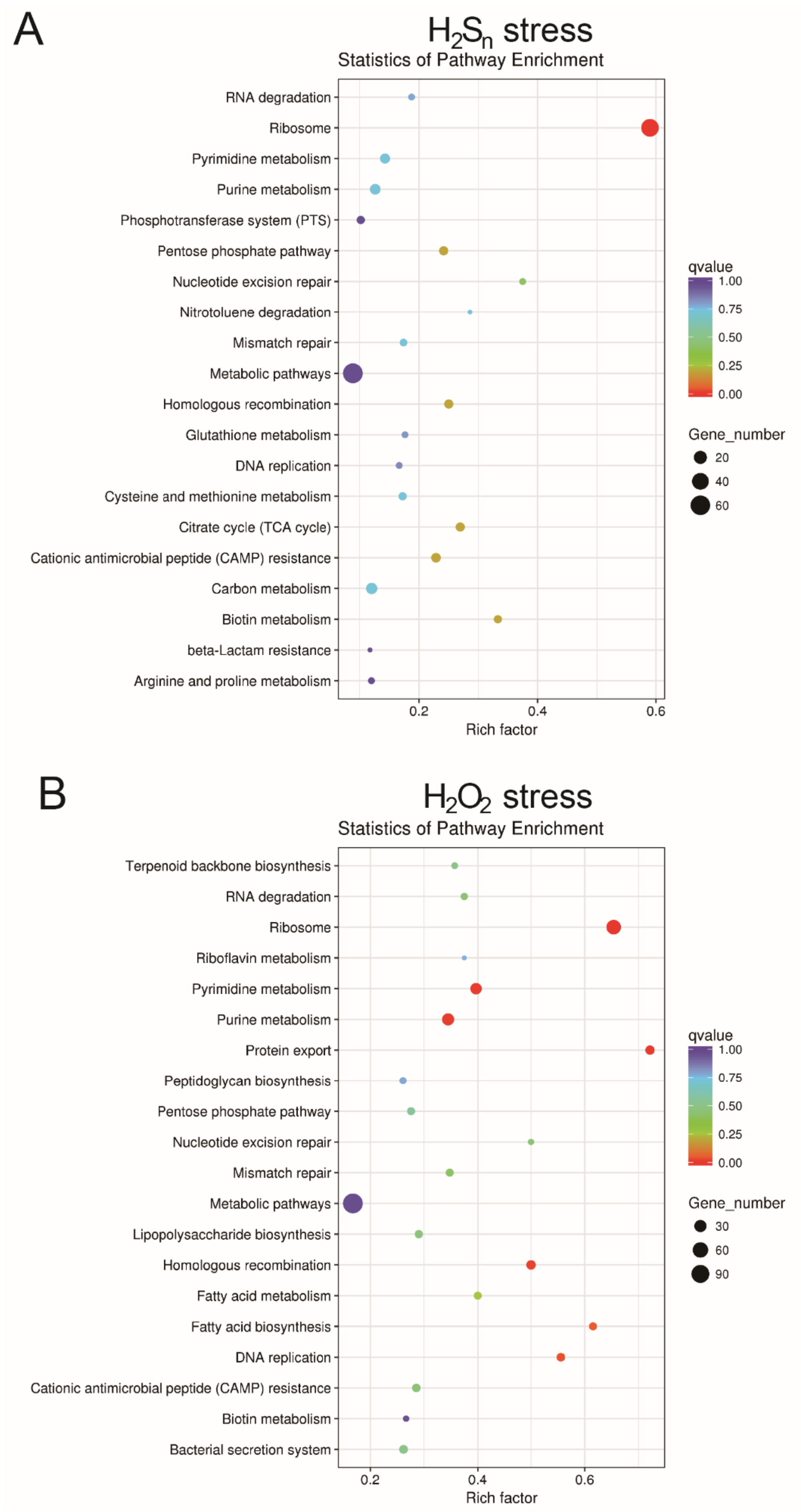


**Figure S7. KEGG metabolism pathway classification of upregulated genes in $H_2S_n$ and $H_2O_2$ stressed *E. coli*.**

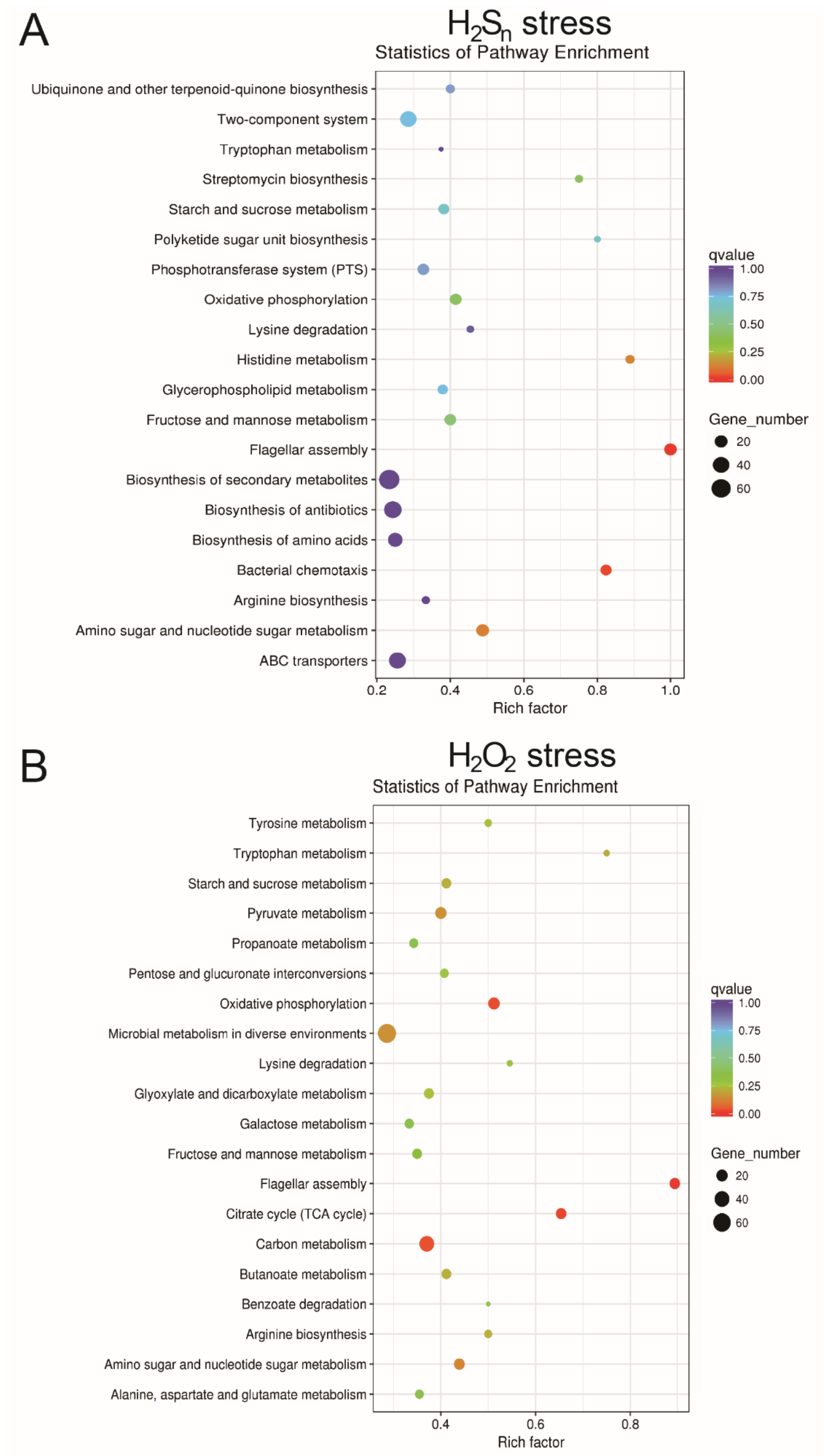


**Figure S8. KEGG metabolism pathway classification of downregulated genes in $H_2S_n$ and $H_2O_2$ stressed *E. coli*.**

**Table S1. The distribution of OxyRs in sequenced bacterial genomes in class level**

| Classes name | Counts |
|---|---|
| Gammaproteobacteria | 2432 |
| Betaproteobacteria | 887 |
| Alphaproteobacteria | 478 |
| Corynebacteriales | 287 |
| Flavobacteriia | 130 |
| Streptomycetales | 67 |
| Bacteroidia | 63 |
| Propionibacteriales | 26 |
| Cytophagia | 24 |
| Deltaproteobacteria | 15 |
| Sphingobacteriia | 13 |
| Deinococci | 11 |
| Oligoflexia | 7 |
| Chitinophagia | 7 |
| Bifidobacteriales | 5 |
| Bacteroidetes Order II. Incertae sedis | 5 |
| Planctomycetia | 5 |
| Actinomycetales | 5 |
| Proteobacteria | 4 |
| Nitrospirales | 3 |
| Micrococcales | 3 |
| Frankiales | 3 |
| Saprospiria | 2 |
| Spirochaetales | 2 |
| Streptosporangiales | 2 |
| Leptospirales | 2 |
| Ignavibacteria | 2 |
| Acidobacteriales | 1 |
| Actinobacteria incertae sedis | 1 |
| Opitutae | 1 |
| Pseudonocardiales | 1 |
| **Total** | **4494** |

### Table S2. Strains and plasmids used in this study

| Entry | Strain/plasmid | Characteristic/description |
|---|---|---|
| | ***Escherichia coli* strains** | |
| 1 | DH5α | supE44, ΔlacU169 (φ80lacZΔM15), hsdR17, recA1, endA1, gyrA96,thi-1, relA1. For cloning and plasmid construction |
| 2 | BL21(DE3) | F-ompT hsdSB (rB-mB-) gal ( λ1857 ind1 Sam7 nin5 lacUV5 T7gene1) dcm. |
| 3 | BL21(DE3) *ΔoxyR* | BL21(DE3) with *oxyR deletion* |
| | ***Plasmids*** | |
| 4 | pTrcHis2A | Amp, Invitrogen. |
| 5 | pTrchis2A-$P_{katG}$-*mkate*-$P_{lacI}$-*oxyR* | *katG* promoter activity reporter with OxyR |
| 6 | pTrchis2A-$P_{grxA}$-*mkate*-$P_{lacI}$-*oxyR* | *grxA* promoter activity reporter with OxyR |
| 7 | pTrchis2A-$P_{trxC}$-*mkate*-$P_{lacI}$-*oxyR* | *trxC* promoter activity reporter with OxyR |
| 8 | pTrchis2A-$P_{katG}$-*mkate*-$P_{lacI}$-$oxyR_{C199S}$ | *katG* promoter activity reporter with $OxyR_{C199S}$ |
| 9 | pTrchis2A-$P_{katG}$-*mkate*-$P_{lacI}$-$oxyR_{C208S}$ | *katG* promoter activity reporter with $OxyR_{C208S}$ |
| 10 | pTrchis2A-$P_{katG}$-*mkate*-$P_{lacI}$-$oxyR_{C199S, C208S}$ | *katG* promoter activity reporter with $OxyR_{C199S, C208S}$ |
| 11 | pTrchis2A-$P_{grxA}$-*mkate*-$P_{lacI}$-$oxyR_{C199S}$ | *grxA* promoter activity reporter with $OxyR_{C199S}$ |
| 12 | pTrchis2A-$P_{grxA}$-*mkate*-$P_{lacI}$-$oxyR_{C208S}$ | *grxA* promoter activity reporter with $OxyR_{C208S}$ |
| 13 | pTrchis2A-$P_{grxA}$-*mkate*-$P_{lacI}$-$oxyR_{C199S, C208S}$ | *grxA* promoter activity reporter with $OxyR_{C199S, C208S}$ |
| 14 | pTrchis2A-$P_{trxC}$-*mkate*-$P_{lacI}$-$oxyR_{C199S}$ | *trxC* promoter activity reporter with $OxyR_{C199S}$ |
| 15 | pTrchis2A-$P_{trxC}$-*mkate*-$P_{lacI}$-$oxyR_{C208S}$ | *trxC* promoter activity reporter with $OxyR_{C208S}$ |
| 16 | pTrchis2A-$P_{trxC}$-*mkate*-$P_{lacI}$-$oxyR_{C199S, C208S}$ | *trxC* promoter activity reporter with $OxyR_{C199S, C208S}$ |
| 17 | pTrchis2A-$P_{katG}$-*mkate* | *katG* promoter activity reporter |
| 18 | pTrchis2A-$P_{grxA}$-*mkate* | *grxA* promoter activity reporter |
| 19 | pTrchis2A-$P_{trxC}$-*mkate* | *trxC* promoter activity reporter |
| 20 | pCL1920 | SPC, low copy plasmid |
| 21 | pCL1920-oxyR native promoter-oxyR | For complement *oxyR* to *ΔoxyR* stain |
| 22 | pTrchis2A-$P_{lacI}$ -*cstR*–$P_{op12}$-*mkate* | CstR-based reporter for detecting intracellular polysulfides |
| 23 | pTrchis2A-$P_{lacI}$-*cstR-mkate-trxA* | CstR-based reporter with *trxA* gene |
| 24 | pTrchis2A-$P_{lacI}$-*cstR-mkate-trxA-trxB*[a] | CstR-based reporter with *trxA* and *trxB* gene |
| 25 | pTrchis2A-$P_{lacI}$-*cstR-mkate-grxB* | CstR-based reporter with *grxB* gene |
| 26 | pTrchis2A-$P_{lacI}$-*cstR-mkate-grxC* | CstR-based reporter with *grxC* gene |

| | | |
|---|---|---|
| 27 | pTrchis2A-$P_{lacI}$-*cstR-mkate-grxD* | CstR-based reporter with *grxD* gene |
| 28 | pBBR1mcs2 | Kmr, broad host range vector |
| 29 | pBBR1mcs2-*katG* | Overexpression of *katG* with *lacI* promoter |
| 30 | pBBR1mcs2-*grxA* | Overexpression of *grxA* with *lacI* promoter |
| 31 | pBBR1mcs2-*trxC* | Overexpression of *trxC* with *lacI* promoter |
| 32 | pET30a | Kmr, expression vector |
| 33 | pET30-*oxyR* | Expression and purification of OxyR with a C terminal His-tag |
| 34 | pET30-$oxyR_{C199S}$ | Expression and purification of $OxyR_{C199S}$ mutant with a C terminal His-tag |
| 35 | pET30-$oxyR_{C208S}$ | Expression and purification of $OxyR_{C208S}$ mutant with C terminal his-tag |
| 36 | pET30-$oxyR_{C199S, C208S}$ | Expression and purification of $OxyR_{C199S, C208S}$ double mutant with C terminal his-tag |

[a] An rbs sequence (ggaaggagattaact) was inserted before *trxA* and *trxB*.
